# Folding latency of fluorescent proteins affects the mitochondrial localization of fusion proteins

**DOI:** 10.1101/768432

**Authors:** Sayaka Kashiwagi, Yoichiro Fujioka, Aya O. Satoh, Aiko Yoshida, Mari Fujioka, Prabha Nepal, Atsushi Tsuzuki, Ozora Aoki, Sarad Paudel, Hitoshi Sasajima, Yusuke Ohba

**Author notes:** **Address correspondence to**: Yusuke Ohba, Department of Cell Physiology, Faculty of Medicine and Graduate School of Medicine, Hokkaido University, N15W7, Kita-ku, Sapporo 060-8638, Japan. **Author contributions:** Conceptualization, Y.F. and Y.O.; investigation, S.K., Y.F., A.O.Y, A.Y., M.F., P.N., A.T., O.A., S.P., and H.S.; writing, S.K. and Y.O.; funding acquisition, Y.F. and Y.O.; supervision, Y.O.

## Abstract

The discovery of fluorescent proteins (FPs) has revolutionized cell biology. The fusion of targeting sequences to FPs enables the investigation of cellular organelles and their dynamics; however, occasionally, such fluorescent fusion proteins (FFPs) exhibit behavior different from that of the native proteins. Here, we constructed a color pallet comprising different organelle markers and found that FFPs targeted to the mitochondria were mislocalized when fused to certain types of FPs. Such FPs included several variants of *Aequorea victoria* green FP (aqGFP) and a monomeric variant of the red FP. Because the FFPs that are mislocalized include FPs with faster maturing or folding mutations, the increase in the maturation rate is likely to prevent their expected localization. Indeed, when we reintroduced amino acid substitutions so that the FP sequences were equivalent to that of wild-type aqGFP, FFP localization to the mitochondria was significantly enhanced. Moreover, similar amino acid substitutions improved the localization of mitochondria-targeted pHluorin, which is a pH-sensitive variant of GFP, and its capability to monitor pH changes in the mitochondrial matrix. Our findings demonstrate the importance of selecting FPs that maximize FFP function.

## INTRODUCTION

Eukaryotic cells contain highly developed membrane-bound organelles with unique morphologies and functions. Each organelle dynamically alters its size, abundance, and intracellular localization in response to a variety of extracellular and intracellular signals. Such organelle dynamics play an important role in cell physiology and pathophysiology. For example, mitochondria are highly mobile structures and continuously change their number, mass, and shape through fusion and fission to maintain functional mitochondria (Archer, 2013; Youle and van der Bliek, 2012). By using quantitative live-cell imaging, it was demonstrated that mitochondria transiently form an enormous, hyperfused network in G1-S phase to enhance their ATP production capacity, thereby playing a regulatory role in G1-S transition (Mitra *et al.*, 2009). Thus, the observation of organelle dynamics is mandatory for a better understanding of biological phenomena at the cellular level.

The morphological examination of organelles had been performed exclusively with electron microscopy until the middle of the last century (Dalton, 1951; Scotuzzi *et al.*, 2017; Seno and Yoshizawa, 1960). This technique allows for the simultaneous observation of all organelles at nanometer resolution; however, live-cell imaging cannot be performed due to the need for fixation (van Zutphen and van der Klei, 2011). The identification of proteins that are specifically localized in a certain organelle (organelle markers) has opened a window for optical microscopy to visualize specific organelles. For instance, the development of specific antibodies against organelle marker proteins has enabled us to paint organelles with colors by an immunofluorescence technique (Johmann and Gorovsky, 1976; Lee *et al.*, 1989; Ralston, 1993). Nevertheless, the fixation process that is required at the first step of this method stops the clock in specimens, thereby discouraging the visualization of organelle dynamics in living cells. Chemically engineered fluorescent dyes targeting specific organelles are available; however, only a limited number of dyes can be used in live-cell imaging because hydrophilic compounds require modifications to break through an aliphatic barrier, the plasma membrane.

Fluorescent proteins (FPs) are powerful tools for live-cell imaging and visualizing organelle dynamics. The introduction of a gene encoding a FP is a simple way to introduce an intrinsic fluorescent dye into living cells because it requires no additional enzymes or cofactors for fluorophore formation (Reid and Flynn, 1997). Because of this advantageous feature, FPs have shed light on molecular dynamics in living cells and even cellular dynamics in living organisms through the development of biosensors to monitor molecular or cellular behaviors (Day and Davidson, 2009; Timpson *et al.*, 2011). Since the first discovery of green fluorescent protein (GFP) in the jellyfish *Aequorea victoria* (Shimomura *et al.*, 1962), the properties of FPs themselves have also evolved through genetic engineering. Such improvements include an extended color pallet (Heim *et al.*, 1994; Ormö *et al.*, 1996), increased fluorescence quantum yield (Bajar *et al.*, 2016; Heim *et al.*, 1995), enhanced folding efficiency at 37ºC (Nagai *et al.*, 2002; Pédelacq *et al.*, 2006), and improved photostability (Griesbeck *et al.*, 2001; Lam *et al.*, 2012; Mena *et al.*, 2006). Despite this series of radical improvements, no versatile FPs have become available thus far. Thus, it is necessary to carefully consider the properties of each FP and the possible unfavorable effects of FP expression in cells.

A good example of this is FP-based organelle markers. Fluorescent fusion proteins (FFPs), which combine FPs and proteins (either full length or portions that include organelle-targeting sequences), are widely used as genetically encoded organelle markers. Such FFPs enable us to visualize time-dependent changes in morphology, subcellular localization, and function in organelles of interest (Chudakov *et al.*, 2010; Rizzuto *et al.*, 1995; Takeuchi and Ozawa, 2007). For instance, the real-time tracking of early, late, and recycling endosomes with FFPs revealed that early endosomes are comprised of two dynamically distinct populations, and such differences are crucial for determining the fate of endocytic cargos in terms of undergoing degradation or recycling (Lakadamyali *et al.*, 2006). However, FFPs do not always exhibit the expected localization. Such mislocalization of FFPs, which might be accounted for by an attribute of either or both the organelle-targeting protein or the FP, is a possible cause of data misinterpretation. For example, the mislocalization in the cytosol of a mitochondria-targeted fluorescent cAMP sensor resulted in “false-positive” findings for changes in mitochondrial cAMP concentration, as it appeared that the cAMP concentration was affected by changes in the cytosol even though it was expected to be unchanged (DiPilato *et al.*, 2004). In fact, an improvement in mitochondrial targeting by fusing repetitive mitochondrial-targeting sequences produced no significant change in the mitochondrial cAMP concentration under the same conditions (Di Benedetto *et al.*, 2013). In addition, the expression of FFPs occasionally affects organelle morphology. In the case of endoplasmic reticulum (ER) markers, the oligomerization of FFPs with dimerizing or tetramerizing FPs resulted in the transformation of an organized smooth ER into a tightly stacked ER (Snapp *et al.*, 2003). Therefore, it is critical to carefully examine whether the developed FFPs indeed localize to the intended organelle and whether they have any adverse effects on organelle morphology or function.

Here, we synthetically constructed a series of FFPs that consist of different organelle-targeting proteins/peptides and a variety of spectral variants of FPs. Among them, we found that mitochondrial matrix markers in combination with FPs with fast maturation rates were mislocalized, as a substantial fraction of such FFPs was observed not in the mitochondria but in the cytoplasm. The cytosolic fraction was significantly reduced by the reintroduction of a mutation that restored the maturation rate to normal. Given that the acceleration of the maturation process is generally thought to be conducive to the performance of FPs, our results suggest that the selection of FPs during the development of FFPs, particularly mitochondrial matrix markers, should be made more carefully.

## MATERIALS AND METHODS

### Reagents and antibodies

MitoTracker™ Red CMXRos and Hoechst 33342 were obtained from Thermo Fisher Scientific (Carlsbad, CA, USA). FCCP and TMRM were purchased from Cayman Chemical (Ann Arbor, MI, USA) and Thermo Fisher Scientific, respectively. An anti-GFP polyclonal antibody (598) and horseradish peroxidase-conjugated anti-rabbit IgG were obtained from Medical and Biological Laboratories (MBL, Nagoya, Japan) and Jackson ImmunoResearch (West Grove, PA, USA), respectively.

### Cell culture

HEK293T (CRL011268), HeLa (CCL-2), Cos-1 (CRL-1650), A431 (CRL-1555), and A549 (CCL-185) cells were obtained from the American Type Culture Collection (Manassas, VA, USA). These cells were cultured under a 5% CO_2_ humidified atmosphere at 37°C in Dulbecco’s modified Eagle medium (DMEM, Sigma-Aldrich, St. Louis, MO, USA) supplemented with 10% fetal bovine serum (Thermo Fisher Scientific). The expression vectors were transfected into HEK293T, HeLa, and Cos-1 cells with “Max” polyethylenimine (Polysciences, Warrington, PA, USA) according to the manufacturer’s recommendations.

### Plasmids

mPlum-Lifeact-7 (Addgene plasmid # 54679) and mCherry-Sec61 β (Addgene plasmid # 49155) were obtained from Drs. Michael Davidson and Gia Voeltz, respectively, via Addgene (Watertown, MA, USA). The expression vectors for Rab5 and Rab7 were described previously (Fujioka *et al.*, 2011). cDNA for each organelle localization sequence was generated by PCR with the following primers: mito_F and mito_R, Tom20_F and Tom 20_R, EEA1_F and EEA1_R, Rab5_F and Rab5_R, Rab7_F and Rab7_R, Rab11_F and Rab11_R, Sec61_F and Sec61_R, Lifeact_F and Lifeact_R, Golgi_F and Golgi_R, Tubulin_F and Tubulin_R, H2B_F and H2B_R, SDHB_F and SDHB_R, and UQCR11_F and UQCR11_R. These sequences were then subcloned into the *Xho*I/*Not*I sites of the pFX-CAAX vectors, which contains human cytomegalovirus (CMV) promoter (Fujioka *et al.*, 2019).

The coding sequences of each fluorescent protein were amplified by PCR with the following primers: EGFP_F and CFP_R, mCherry_F and mCherry_R, iRFP_F and iRFP_R, MiCy_F and MiCy_R, mKate2_F and mKate2_R, TFP650_F and TFP650_R, pHluorin_F and pHluorin_R, sfGFP_F and sfGFP_R, and hdKeima_F and hdKeima_R. These sequences were then subcloned into the *Eco*RI/*Bgl*II, *Eco*RI/*Xho*I, or *Not*I/*Bgl*II sites of the pFX vectors.

The coding sequence of AcGFP1 was amplified by PCR with the AcGFP_F and AcGFP_R primers and subcloned into the *Age*I/*Not*I sites of pDsRed2-Mito (Clontech). The coding sequence of Venus (cDNA was obtained from Dr. Atsushi Miyawaki; Nagai *et al.*, 2002) with four mutations (E116Q/G117D/D118G/T119C, RFPized Venus, DsVenus) was obtained by PCR-based mutagenesis with the following primers: EGFP_F and QDGC_R and QDGC_F and CFP_R. After gel extraction, the two products were mixed and subjected to a 2^nd^ PCR to obtain the full-length DsVenus sequence with the primers EGFP_F and CFP_R. To generate the sequence encoding the A163V and/or G175S mutants of pHluorin, PCR-based mutagenesis with a QuickChange Site-Directed Mutagenesis Kit (Agilent Technologies, Santa Clara, CA, USA) was performed with the primers A163V_F and/or G175S_F. The primers used in this study are listed in Table S1.

### Fluorescence Microscopy

Cells were imaged on either an IX-83 or IX-81 inverted microscope (Olympus, Tokyo, Japan), both of which were equipped with a BioPoint MAC 6000 filter and shutter control unit (Ludl Electronic Products, Hawthorne, NY, USA), an automated XY-stage (Chuo Precision Industrial, Tokyo, Japan), and a SOLA Light Engine (Lumencor, Beaventon, OR, USA) as an illumination source. UPlanSApo 60×/1.35 oil objective lenses and UPlanSApo 10×/0.40 objective lenses were used. The following excitation and emission filters were used in this study: FF01-387/11-25 and FF02-447/60-25 (Semrock, Rochester, NY, USA) for Sirius and Hoechst 33342; FF02-438/24 and FF01-483/32 (Semrock) for CFP and its derivatives; BP470-490 and BP510-550 (Olympus) for the MiCy, GFP, and GFP derivatives; FF01-500/24-25 and FF01-542/27 (Semrock) for YFP and its derivatives; BP520-550 and BA580IF (Olympus) for DsRed, mCherry, mKate2, eqFP650, MitoTracker™, and TMRM; and FF02-628/40 and FF01-692/40 (Semrock) for iRFP. Confocal images were acquired with an sDISK spinning disk unit (Andor Technology, Belfast, UK) and a Rolera EM-C^2^ electron multiplying cooled charge-coupled device camera (QImaging, Surrey, BC, Canada), whereas the epifluorescence images were acquired with a Cool SNAP MYO cooled charge-coupled device camera (Photometrics, Tucson, AZ, USA). MetaMorph software (Molecular Devices, CA, USA) was used for the control of the microscope and the peripheral equipment. For live-cell imaging, the atmosphere was maintained at 37°C with a Chamlide incubator system (Live cells instrument, Seoul, Korea) for both microscopes.

For live cell imaging, cells plated on collagen-coated 35-mm-diameter glass-bottom dishes (AGC techno glass, Shizuoka, Japan) were transfected with expression vectors for FFPs as indicated in the figure legends. Unless otherwise noted, the cells were transferred into phenol red-free DMEM/F12 (Thermo Fisher Scientific) 24 h after transfection and were imaged with the microscopes. For some experiments, the cells were labeled with MitoTracker™ Red CMXRos (12.5 nM) for 10 min at 37°C (for the determination of mitochondrial morphology and fluorescent intensity) or TMRM (50 nM) and Hoechst 33342 (1 µg/ml) for 30 min at 37ºC (for the measurement of mitochondrial pH) before imaging. Alternatively, cells plated on collagen-coated 96-well glass-bottom plates (AGC techno glass) were transfected with expression vectors. After 24 h, the cells were fixed in 3% paraformaldehyde for 15 min at room temperature, transferred into PBS, and then imaged.

### Quantification of FFP Localization and Mitochondrial Fragmentation

Mitochondrial masked images were generated based on images obtained through the MitoTracker channel as described previously with slight modifications (Iannetti *et al.*, 2016). In brief, the background-corrected MitoTracker image was subjected to the ‘rolling ball’ collection algorithm. The resulting image was processed with a Mexican Hat filter to extract the particle objects, followed by noise reduction with a Median filter. A mitochondrial binary image was obtained by the ImageJ plugins “Auto Threshold” and ‘Remove Outliers.’ Fluorescence intensity in the mitochondrial area in the masked image was recorded for the determination of the mitochondrial intensity. The fluorescence intensity of the whole cell was measured within the area that used for the determination according to the DIC images. The ratio of the mitochondrial intensity to the whole cell intensity was calculated to evaluate the extent of the mitochondrial localization of the FFPs.

For the analysis of mitochondrial fragmentation, according to the mitochondrial mask images obtained as described above, the number of mitochondrial particles × 10,000/total area of the mitochondria (in pixels) was calculated and plotted as the mitochondrial fragmentation count [AU] as described previously (Rehman *et al.*, 2012).

### In Vitro pH Titration

Total cell lysates were prepared as described in the immunoblotting section and were mixed with 20 mM citrate phosphate buffer (containing 150 mM NaCl) to obtain a pH value ranging from 4.5 to 8.5. The mixtures (100 μL each) were transferred into a well of a Nunc black polystyrene 96-well Microwell™ plate (Thermo Fisher Scientific). The fluorescence emission at 530 nm ± 15 nm was measured using a SpectraMax i3x multimode microplate reader (Molecular Devices) at an excitation wavelength of 470 nm ± 9 nm. After the subtraction of background fluorescence (from the lysis buffer), the fluorescence intensities were plotted against the pH and fitted to the four-parameter logistic curve (which is mathematically analogous to the Hill equation; DeLean *et al.*, 1978)

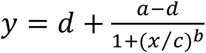

by using the least-squares method. In this equation: *y*, fluorescence intensity; *x*, proton concentration; *a*, the fluorescence intensity when *x* = 0; *d*, the fluorescence intensity when x = ∞ *c*, the proton concentration when *y* = (*a* + *d*)/2; *b*, the steepness of the curve. p*K*_a_ was calculated with the equation p*K*_a_ = - log *c*. The data shown in Figure 6A are the mean ± standard error of the mean (s.e.m.) for the values obtained when the maximum (*d*) and minimum values (*a*) were normalized to 1 and 0, respectively.

### Measurement of Mitochondrial pH

Cos-1 cells expressing mito-pHluorin or mito-*at* pHluorin were pretreated with TMRM and Hoechst 33342 and subjected to time-lapse fluorescence microscopy. At time 0, the cells were exposed to 10 µM FCCP, which is an uncoupler that lowers both the cytosolic and mitochondrial pH (Berezhnov *et al.*, 2016). After background subtraction, the fluorescence intensity within the cells was quantified by using the Multi Wavelength Cell Scoring module in MetaMorph. The weighted average of the fluorescence intensity at each time point normalized to that at time 0 was calculated and plotted.

### Statistical Analyses

The quantitative data are presented as the mean ± s.e.m. of at least three independent experiments (unless indicated otherwise) and were compared by one-way analysis of variance (ANOVA) followed by Dunnett’s post hoc analysis (multiple conditions against a control condition) or one-way analysis of variance (ANOVA) followed by a post hoc Tukey honestly significant difference (HSD) test (among multiple conditions). The time series data sets were compared by multivariate analysis of variance (MANOVA). No statistical methods were used to predetermine the sample size. The studies were performed unblinded.

## RESULTS

### Development of a Color Pallet of Organelle-targeted FFPs

To perform the simultaneous observation of dynamics in multiple organelles in single living cells, we constructed a set of FFPs that were targeted to organelles and fused with a variety of FPs. The organelles and cell architectural features tested include the mitochondria (matrix and outer membrane), endosomes (early, late, and recycling), ER, Golgi apparatus, actin, microtubules, plasma membrane, and nucleus, which were targeted by the mitochondria-targeting sequence of the cytochrome c oxidase subunit VIII (simply called mito hereafter), translocase of outer membrane 20 (TOM20), early endosomal antigen 1 (EEA1), several small GTPases (Rab5, Rab7, and Rab11), Sec61 β, β1,4-galactose transferase (GalT), Lifeact (a *Saccharomyces cerevisiae*-derived peptide), tubulin β-4B chain, the C-terminal hypervariable region of K-Ras 4B (KRasCT), and histone 2B (H2B), respectively (Fujioka *et al.*, 2018; Lakadamyali *et al.*, 2006; Riedl *et al.*, 2008; Shaner *et al.*, 2008; Wu *et al.*, 2016; Zurek *et al.*, 2011). The FPs utilized included Sirius, SECFP, EGFP, Venus, mCherry, and iRFP (Filonov *et al.*, 2011; Nagai *et al.*, 2002; Shaner *et al.*, 2008; Tomosugi *et al.*, 2009). Cos-1 cells (Figures 1 and S1) and HeLa cells (Figure S2) were transfected with expression vectors for these FFPs and observed with confocal microscopy. Most of the FFPs displayed the expected localization in the targeted organelles or structures without any visible abnormality in terms of cell morphology, except that the excess expression of TOM20-iRFP occasionally induced mitochondrial aggregation (Figure S3A). For example, FP-tagged Lifeact, an F-actin marker, clearly depicted a fibrous structure, while the Golgi apparatus-localizing enzyme GalT was localized only to the perinuclear region (Figures 1, Figure S1 and S2). However, some FFPs were also found to be distributed to a compartment different from that of the expected organelle in a manner that was dependent on the utilized FP. For example, whereas EGFP-tagged mito, a common mitochondria marker (De Michele *et al.*, 2014), clearly localized only to the mitochondria, other FFPs exhibited “cytosolic and nuclear leaks” in addition to their expected localization (mitochondria) (Figure 1, left column). Besides the mitochondrial matrix markers, endosome-targeted FFPs showed a similar tendency (Figure S3B).

**Figure 1:**
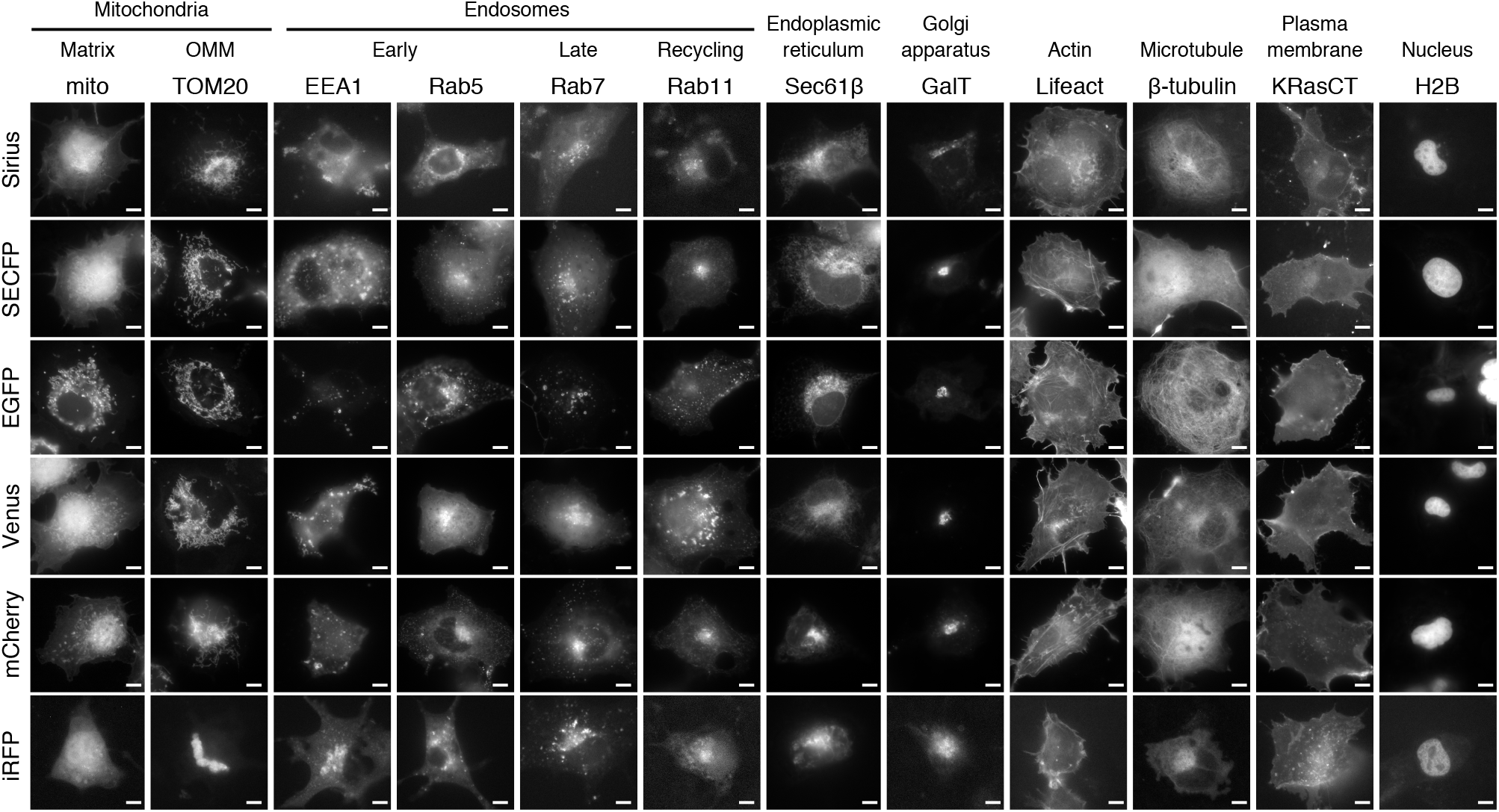
A color pallet of the organelle-targeted FFPs. Cos-1 cells were transfected with expression vectors for FFPs (organelle targeted and FP used are indicated at the top and left, respectively) for 24 h and observed with confocal microscopy. OMM: outer mitochondrial membrane. Representative images are shown. Bar, 10 μm.

### Tagged FP-dependent Localization of Mitochondrial Matrix Markers

To determine whether the mislocalization of FFPs was due to the mitochondrial-targeting sequences or the FPs, alternative mitochondrial matrix markers with mitochondrial proteins other than mito were prepared. We used succinate dehydrogenase [ubiquinone] iron-sulfur subunit (SDHB, a mitochondrial matrix protein) and cytochrome b-c1 complex subunit 10 (UQCR11, a mitochondrial inner membrane protein) because these proteins, when tagged with the GFP variant mCitrine, have been reported to be localized to the mitochondria (Söhnel *et al.*, 2016). Nevertheless, SECFP-tagged SDHB displayed similar cytoplasmic and nuclear mislocalization as that of mito-SECFP. Moreover, to make matters worse, its expression induced severe mitochondrial fragmentation (Figure S4). The localization of UQCR11-SECFP was more mitochondria specific; however, it resulted in mitochondrial swelling and fragmentation (Figure S4). These results together indicated that mitochondrial-targeting sequences might not dictate the mitochondrial localization of FFPs. In addition, the mitochondrial targeting sequences used for developing FFPs should be selected with care because the mitochondrial architecture is likely to be susceptible to the overexpression of component proteins.

We therefore prepared mitochondria matrix markers that consisted of mito and other color variants of FPs to evaluate the effects of different types of FPs on FFP localization. The transfection of pDsRed-mito, a commercially available expression vector for mito tagged with RFP from *Discosoma* sp., into Cos-1 cells resulted in distribution exclusively in the mitochondria (Figure 2). We also prepared expression vectors for mito tagged with MiCy, mKate2, and eqFP650 (Karasawa *et al.*, 2004; Shcherbo *et al.*, 2009; 2010), none of which originate from *Aequorea victoria*, and found that these FFPs also localized preferentially to mitochondria (Figure 2). In contrast, Sirius, SECFP, Venus, and mCherry-tagged mito failed to do so (Figure 2). Such mislocalization of mitochondria-targeted FFPs was also observed in other cell lines besides Cos-1 cells (Figures S5, A and B), indicating that this was caused by the properties of the FFPs but not by the cell context. It was previously reported that similar mislocalization of a mito-tagged biosensor based on the principle of Förster resonance energy transfer (FRET) was observed 24 h after transfection, whereas it was no longer present at 48 h after transfection (Di Benedetto *et al.*, 2013). Therefore, we obtained a series of cell images at 24, 48, and 72 h after transfection, which did not show any significant differences in localization (Figures S5, C and D).

**Figure 2:**
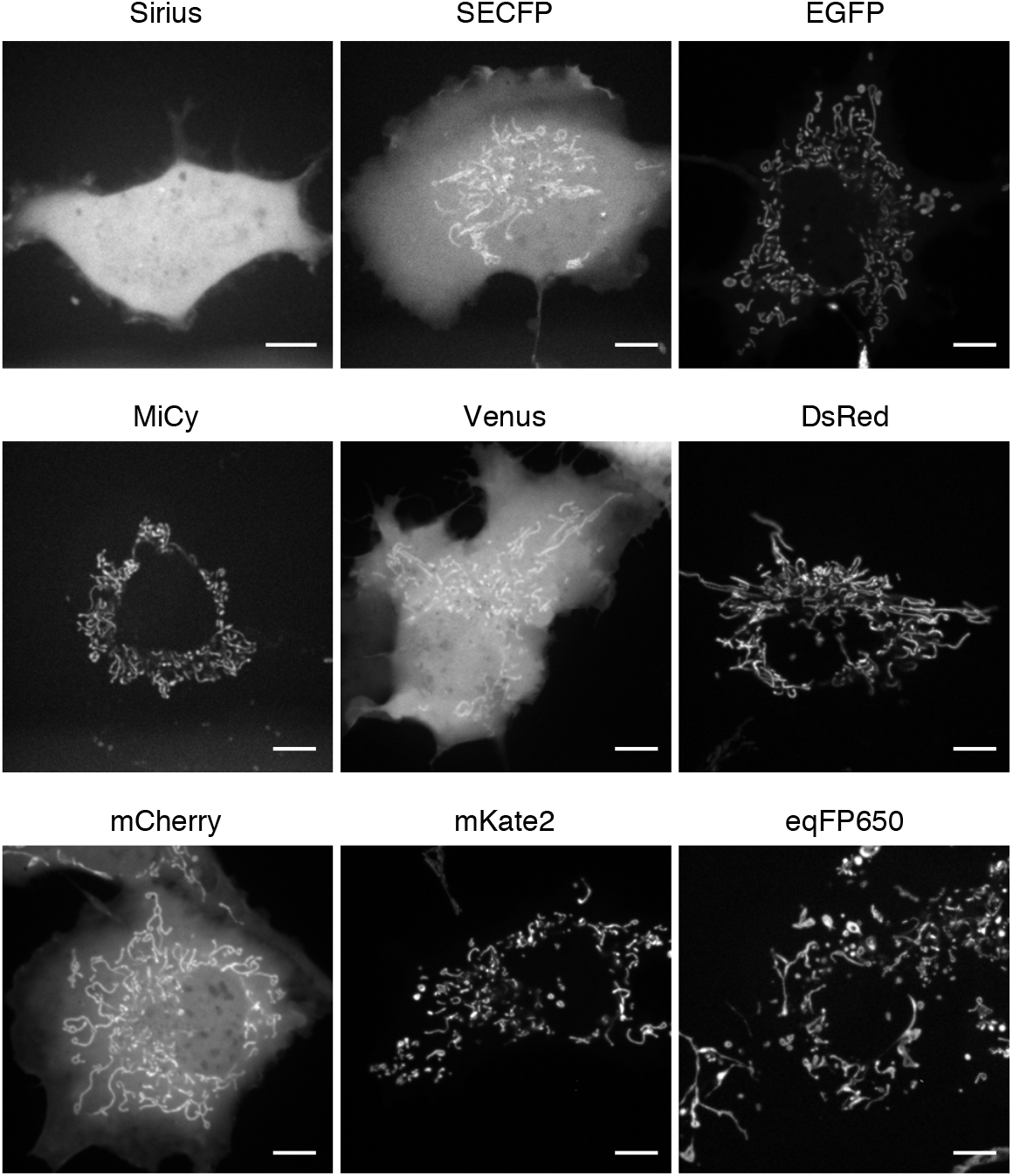
Subcellular localization of mitochondria-targeted FFPs. Cos-1 cells expressing mitochondria-targeted FFPs that contained the FPs indicated at the top were observed with confocal microscopy. Representative images are shown. Bar, 10 μm.

### Construction of the RFPized Venus containing FFP

Next, we examined the properties of the FPs utilized for the mitochondrial markers and noticed that almost all the mislocalized FFPs, except for mito-EGFP, utilized FPs derived from *Aequorea victoria*. In addition, the majority of the FFPs that harbored RFP (DsRed, mKate2, and eqFP650, but not mCherry) was clearly localized only to mitochondria (Figure 2). To examine whether the origin of the FPs affected the mitochondrial localization of the markers, we generated red fluorescent proteinized (RFPized) Venus, into which several amino acid substitutions (E116Q/G117D/D118G/T119C) was introduced (DsVenus) and constructed an expression vector for mito-DsVenus (Figure 3A). However, unexpectedly, mito-DsVenus was still localized in the cytosol and nucleus in addition to mitochondria (Figure 3B). To quantify the extent of the mitochondrial localization of the markers, we calculated the ratios of the fluorescence intensity in the mitochondria to those in the cytosol and nucleus (Figure S3A). Although the mitochondrial localization of mito-DsVenus was likely to be higher than that of mito-Venus, there was no significant difference between the two conditions (Figure 3C). These results indicated that the origin of the FP does not affect the localization of mitochondria-targeted FFPs.

**Figure 3:**
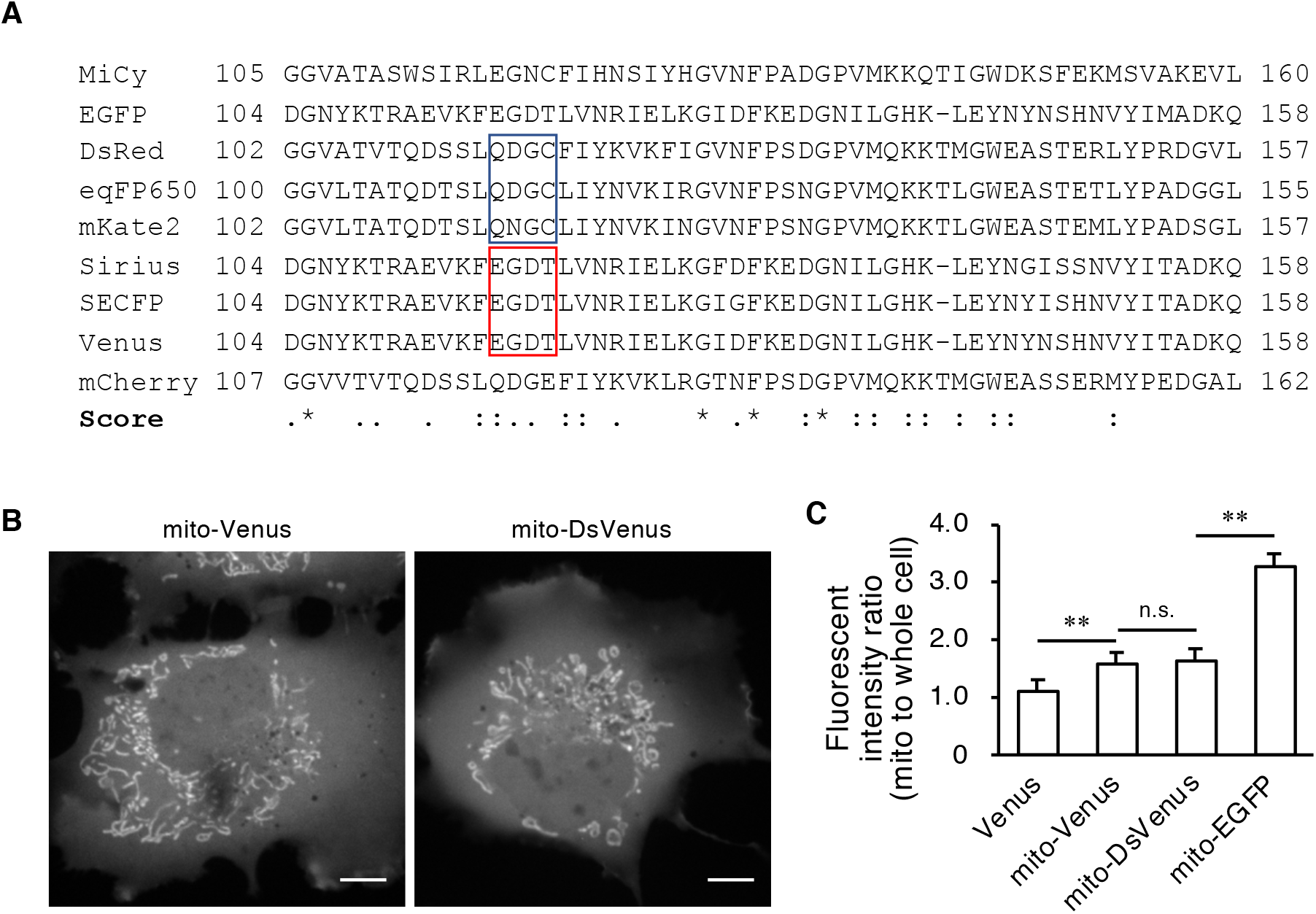
Development of FFPs with RFPized FPs. (**A**) Comparison of the amino acid sequences of the FPs used in this study. Blue and red boxes indicate amino acids conserved in the RFP and GFP groups, respectively, but that are distinct among the groups. “.”, “:”, and “*” indicate low homology, high homology, and identical amino acids, respectively. (**B**) Cos-1 cells expressing the FFPs indicated at the top were observed with confocal microscopy. Representative images are shown. (**C**) Cos-1 cells expressing the FFPs indicated at the bottom were observed with confocal microscopy, and the extent of mitochondrial localization was quantitated and plotted. The data shown represent the mean ± s.e.m. from three independent experiments (*n* > 10). **, *p* < 0.0001, as calculated by one-way ANOVA with a post hoc Tukey HSD test. n.s., not significant.

### Mislocalization of Mitochondria-targeted FFPs Induced by Folding Mutations

We next generated mitochondria-targeted FFPs with two *Aequorea victoria*-derived FPs, roGFP2 (Hanson *et al.*, 2004; hereafter roGFP) and superecliptic pHluorin (Sankaranarayanan *et al.*, 2000; hereafter pHluorin), as well as *Aequorea coerulescens*-derived GFP, AcGFP1 (Gurskaya *et al.*, 2003; hereafter AcGFP). roGFP and pHluorin are widely used as subcellular redox and pH sensors, respectively. It is worth noting that commercially available mito-AcGFP is shown to be localized only to the mitochondria. It was revealed that roGFP- and AcGFP-tagged mito were exclusively localized to the mitochondria (Figure 4A), whereas pHluorin-tagged mito exhibited cytosolic and nuclear leakage, as did SECFP- or Venus-tagged mito. We collectively compared the amino acid sequences of the utilized FPs and found that amino acid substitutions (M153T/V163A/S175G) that promote their protein folding (so-called “folding mutations”; Nagai *et al.*, 2002) had been introduced into the FPs of the mislocalized FFPs (Figures 4B and 4C).

**Figure 4:**
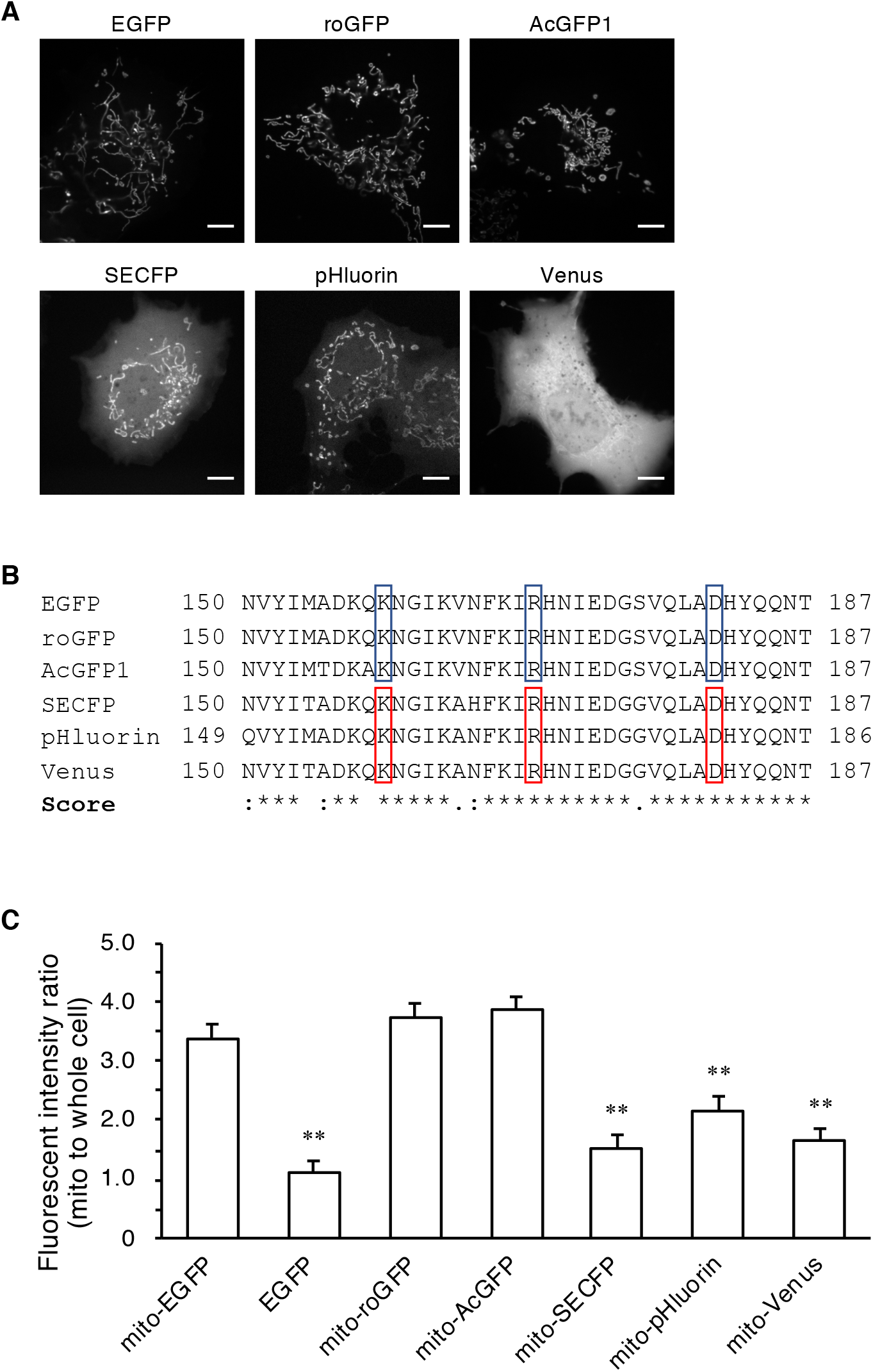
Mislocalization of mitochondria-targeted FFPs with fast folding FPs. (**A**) Cos-1 cells expressing mitochondria-targeted FFPs that harbored the FPs indicated at the top were observed by confocal microscopy. Representative images are shown. Bar, 10 μm. (**B**) Comparison of the amino acid sequences of the FPs. Blue and red boxes indicate the amino acids of the original aqGFP and those in the fast folding mutant, respectively. “.”, “:”, and “*” indicate low homology, high homology, and identical amino acids, respectively. (**C**) Cos-1 cells expressing the FFPs indicated at the bottom were observed with confocal microscopy, and the extent of mitochondrial localization was quantitated and plotted. The data shown represent the mean ± s.e.m. from three independent experiments (n > 10). **, *p* < 0.0001, as calculated by one-way analysis of variance (ANOVA) followed by Dunnett’s post hoc analysis. n.s., not significant versus mito-EGFP. Bar, 10 μm.

### Improvement of the Mitochondrial Localization of FFPs by Reverse Mutation of the FPs

The above results raised the possibility that folding mutations in FPs inhibited the accurate mitochondrial localization of FFPs. To test this hypothesis, we prepared mito-tagged EYFP and mito-tagged Venus and compared their localization. These two FPs have essentially same excitation/emission spectra, but only the latter has a folding mutation. As expected, whereas the fluorescence signal of mito-Venus was mainly observed in the cytosol and nucleus, mito-EYFP was preferentially localized to the mitochondria (Figures 5, A and B). We further evaluated the localization of mito tagged with superfolder GFP (sfGFP corresponds to aqGFP M153T/V163A), which is a fast-folding EGFP variant generated by DNA shuffling (Pédelacq *et al.*, 2006). In contrast to mito-EGFP, mito-sfGFP showed mislocalization in the cytosol and nucleus (Figure 5C). To provide more evidence for this notion, we constructed a pHluorin revertant in which the folding mutations of pHluorin (V163A/S175G) were restored to those of the original aqGFP (V163/A175). The revertant pHluorin-tagged mito was indeed clearly localized to the mitochondria, and the degree of this localization compared to that of the original pHluorin-tagged mito was dependent on the number of restored amino acids (Figures 5, E and F). The double mutant was therefore named *at* pHluorin after the musical term “*a tempo*,” which means “at original speed” or “cancelation of the last speed change.”

**Figure 5:**
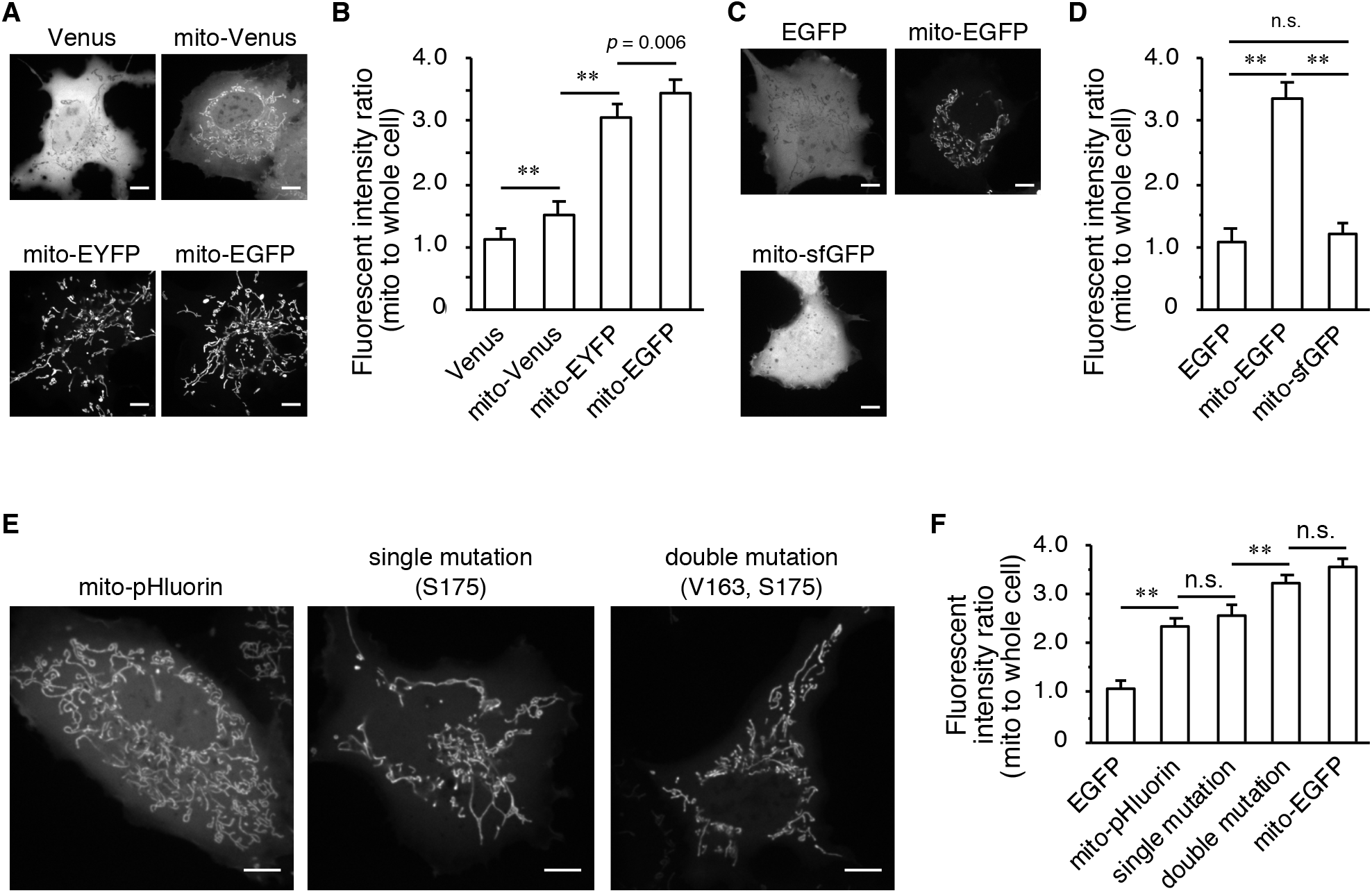
The *at* mutation improved the mitochondrial localization of FFPs. Cos-1 cells expressing the FFPs indicated at the top were observed by confocal microscopy. Representative images are shown (**A**, **C**, **E**). Bar, 10 μm. The extent of mitochondrial localization was quantitated and plotted (**B**, **D**, **F**). The data shown represent the mean ± s.e.m. from three independent experiments (*n* > 10). The *p*-values were calculated by one-way ANOVA with a post hoc Tukey HSD test. **, *p* < 0.0001, n.s., not significant.

### Precise Measurement of Mitochondrial pH by at pHluorin

We finally compared the properties of mitochondria-targeted FFPs with pHluorin and *at* pHluorin in terms of specific measurement of the mitochondrial pH. An in vitro titration assay revealed that the amino acid substitution resulted in no difference in pH sensitivity (Figure 6A). Under these conditions, changes in pH in the mitochondrial matrix induced by carbonyl cyanide p-trifluoromethoxyphenylhydrazone (FCCP) were evaluated. FCCP is a chemical uncoupler that promotes mitochondrial membrane permeability to proton ions and acidifies the mitochondrial matrix (Aw and Jones, 1989). Intriguingly, while the fluorescence intensities of both mito-pHluorin and mito-*at* pHluorin were decreased by FCCP treatment, the extent of the decrease in the intensity of mito-*at* pHluorin was smaller than that of mito-pHluorin (Figure 6B). However, when the changes in mitochondrial pH were separately quantified via mitochondrial masked images, which were generated based on the images of the chemical mitochondrial marker MitoTracker™ Red CMXRos as a guide, it was revealed that mito-*at* pHluorin indeed reported essentially the same dynamics in terms of pH change in mitochondria as those reported by mito-pHluorin (Figure S6A). Given that FCCP was reported to decrease the cytosolic pH more than that of the mitochondrial matrix by increasing plasma membrane permeability to proton ions (Berezhnov *et al.*, 2016), the presence of cytoplasmic mito-pHluorin resulted in the overestimation of the mitochondrial pH. Indeed, the fluorescence intensity of cytosolic pHluorin was decreased by FCCP treatment (Figure S6B). Overall, the preferential localization of mito-*at* pHluorin enables the specific measurement of pH changes in the mitochondrial matrix without the use of other complementary fluorescent dyes. This would be advantageous for multidimensional imaging with multiple FFPs and for flow cytometry analyses.

**Figure 6:**
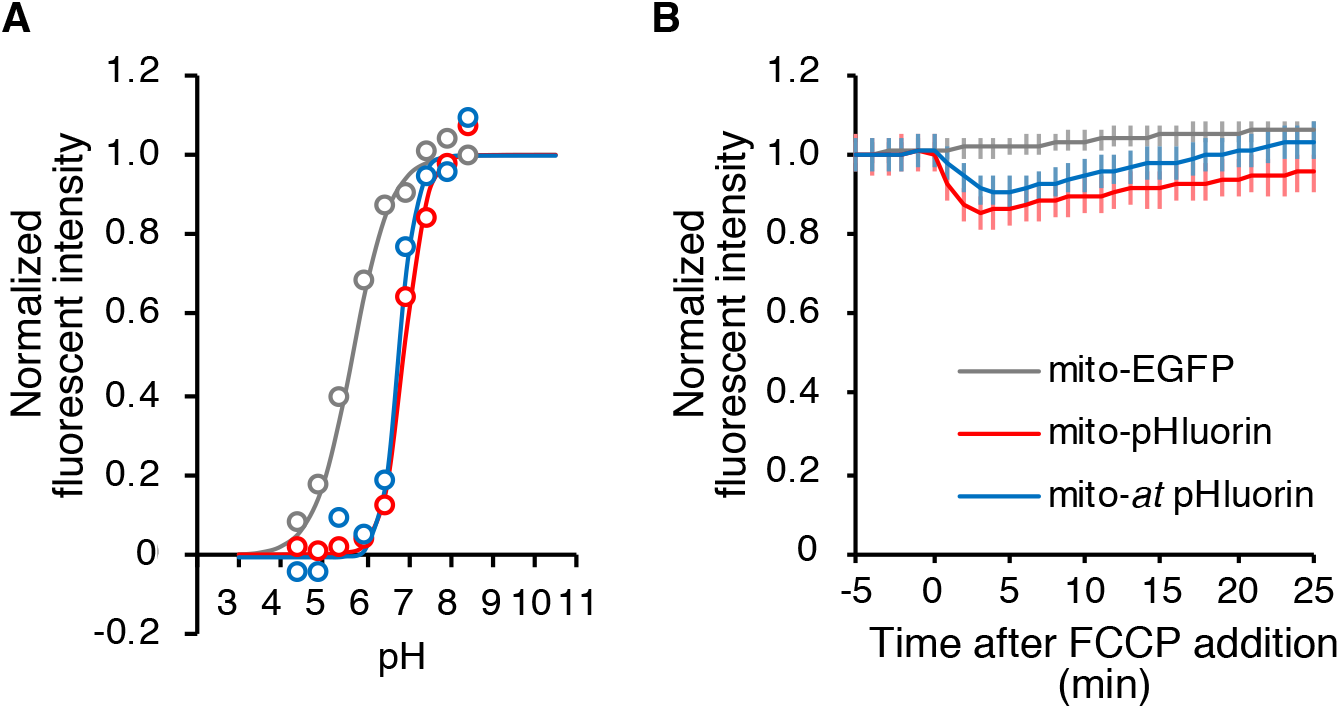
Measurement of mitochondrial pH by FFPs. (**A**) HEK 293T cells expressing either mito-EGFP, mito-pHluorin, or mito-*at* pHluorin were subjected to an in vitro titration assay. The fluorescence intensities at each pH were normalized to the maximum intensity and plotted against the pH. The data were fitted to Hill’s equation, and the fitted curves are also shown. The *p*K_a_ for mito-EGFP, mito-pHluorin, and mito-*at* pHluorin was 5.80, 7.05, and 6.98, respectively. (**B**) Cos-1 cells expressing the proteins indicated at the bottom were subjected to time-lapse fluorescence microscopy. At time 0, the cells were exposed to FCCP. The normalized fluorescence intensities were plotted over time. The data shown are the mean ± s.e.m. from three independent experiments (*n* > 145). *p*-values were calculated by MANOVA. **, *p* < 0.0001, n.s., not significant.

## DISCUSSION

In this study, we constructed more than 80 organelle-targeted FFPs and found that several sets of FFPs localized to organelles other than those expected. In the case of mitochondrial matrix markers, it was demonstrated that mutations that accelerate the folding process of FPs reduced the expected (mitochondrial) localization. Endosomal markers also exhibited a similar tendency. Following the discovery and cloning of GFP and its relatives, FPs have facilitated the routine monitoring of protein and organelle dynamics in living cells and cellular dynamics in whole animal bodies (Chudakov *et al.*, 2010; Day and Davidson, 2009). When tagging a protein of interest with FPs, FPs are usually appended to the amino and/or carboxyl terminus or even added within the sequence of a protein. The choice of the portion to which FPs are attached can be determined according to the reported properties of the protein of interest; however, in most cases, there is no rational basis for the design of FFPs and a trial-and-error strategy must be used. Because of the size of a fluorescent protein, the resulting perturbations to the overall folding and function of the protein of interest can be significant. When designing an FFP, linkers can be used to join the two sequences. This strategy helps to overcome the aforementioned challenges of protein folding and function, but the sequence itself is also critical for the proper functioning of tagged proteins. In addition to the above issues (the site and method used to add FPs), we propose that the properties of the FP itself can determine FFP function, particularly in the case of mitochondrial and endosomal markers.

Major mitochondrial proteins are synthesized in the cytosol and transported to the mitochondria. To be localized within mitochondria, the precursors need to be imported across one or both of the mitochondrial membranes through a pore formed by the TOM complex (Chacinska *et al.*, 2009). Given the diameters of pores, which range from 13 Å (Tim23) to 20 Å (Tom40), only unfolded proteins can pass through them (Pfanner and Truscott, 2002). It is generally assumed that mitochondrial preproteins are usually unfolded or loosely folded in the cytosol and guided by cytosolic targeting factors or chaperones to the mitochondria. Furthermore, it has also been reported that several preproteins contain stably folded domains in the cytosol (Bömer *et al.*, 1997; Wienhues *et al.*, 1991). For such proteins, the mitochondrial import machinery actively unfolds and imports the preprotein. Indeed, the importing efficiency of a protein is correlated with the ease of its unfolding and resistance to mechanical unfolding of the folded proteins (Eilers and Schatz, 1986; Sato *et al.*, 2005; Wilcox *et al.*, 2005). In the case of mitochondria-targeted FFPs with aqGFP, we clearly demonstrated that the fast folding of FPs prevented their proper localization. It is noteworthy that only mCherry is found to be mislocalized among the RFPs tested. Given that mCherry horbors the mutations V7I and M182K, which promote the protein folding (half time for maturation of DsRed and mCherry are ~10h and 15 min, respectively) (Shaner *et al.*, 2008), the relationship between folding latency and mitochondria targeting might serve as a common future for different FPs. It might be possible that once fast-folding FPs are folded, they become stable in the folded state and are resistant to unfolding by the mitochondrial protein import machinery. Indeed, it has been reported that GFP39N (F99S/M153T/V163A) displays higher thermostability than wild-type aqGFP (Aliye *et al.*, 2015). Another possibility is that folded FPs prevent the binding of FFPs to mitochondrial-targeting factors or chaperones in the cytosol, which guide them to the mitochondrial surface. For FFPs to be incorporated into mitochondria, the FPs within FFPs must be in an unfolded state. Given that mito-DsRed, which is more stable than EGFP when folded (Verkhusha *et al.*, 2003), and mito-EGFP were preferentially localized to the mitochondria, the latter possibility might be more plausible. A systematic examination of the mechanical unfolding kinetics of FPs by atomic-force microscopy will be of great help in clarifying this issue (Perez-Jimenez *et al.*, 2006).

Our observations demonstrated that mutations that accelerate the folding process of FPs disturb the mitochondrial localization of FFPs. There are a great number of FP variants that have been developed with the aim of improving brightness (via increased quantum yield and folding efficacy at 37ºC), changing the excitation and emission spectra, and decreasing the chemical and photochemical sensitivity (Day and Davidson, 2009; Tsien, 1998). Among them, those with mutations that improve the folding efficiency of FP at 37ºC (often called “folding” mutations; (Fukuda *et al.*, 2000) exhibit enhanced brightness due to an increased proportion of mature, light-emitting FP molecules. Therefore, these mutations are generally considered to be favorable, which has encouraged the construction of new FFPs and the substitution of classical FPs with improved FPs. These results may lead to the questioning of this trend because fast-folding FP variants hamper the correct localization of FFPs to the mitochondrial matrix, and a reverse mutation causes them to localize specifically to the mitochondria. A similar tendency was also observed for the endosomal markers.

In conclusion, we hereby suggest that slow-folding FPs might occasionally be suitable for FFPs targeting certain organelles, including the mitochondria and endosome. An increase in the extent of specific localization to the targeted organelle will improve the signal to noise ratio, which will improve the detection of not only organelle morphology but also organelle-specific changes in ion concentration, signaling, and protein-protein interactions with FFPs. This will definitely improve the elucidation of cell physiological functions in detail.

## Abbreviations used

aq: *Aequorea victoria*
Ds: *Discosoma* sea anemones
EEA1: early endosomal antigen 1
ER: endoplasmic reticulum
FCCP: p-trifluoromethoxyphenylhydrazone
FFP: fluorescent fusion proteins
FP: fluorescent proteins
FRET: Förster resonance energy transfer
GalT: β1,4-galactose transferase
GFP: green fluorescent protein
H2B: histone 2B
KRasCT: C-terminal hypervariable region of K-Ras 4B
RFPized: red fluorescence-proteinized
sfGFP: superfolder GFP
TOM: translocase of outer membrane

## ACKNOWLEDGEMENTS

We thank A. Miyawaki for providing the Venus cDNA and A. Kikuchi for technical assistance. This work was supported in part by Grants-in-Aid from the Ministry of Education, Culture, Sports, Science and Technology of Japan (#18H04850, #19H05411, and #19H04823) and the Japan Society for the Promotion of Science (#17H04016, 19K22506, #16H06227, 17J07984, and 18KK0196) as well as by grants from the Canon Foundation, the. Akiyama Life Science Foundation, and the Konica Minolta Science and Technology Foundation.

---

**Figure S1:**
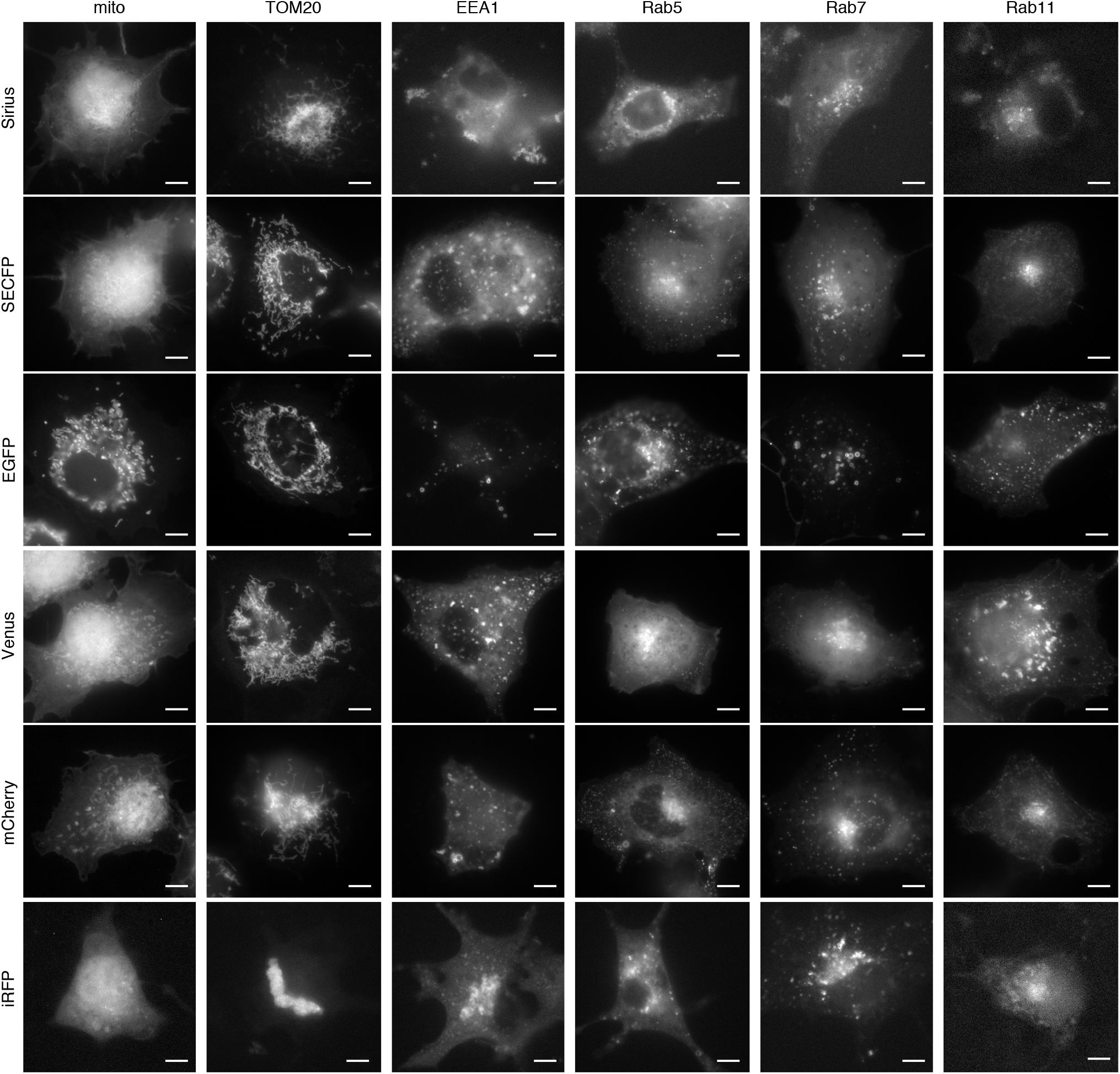
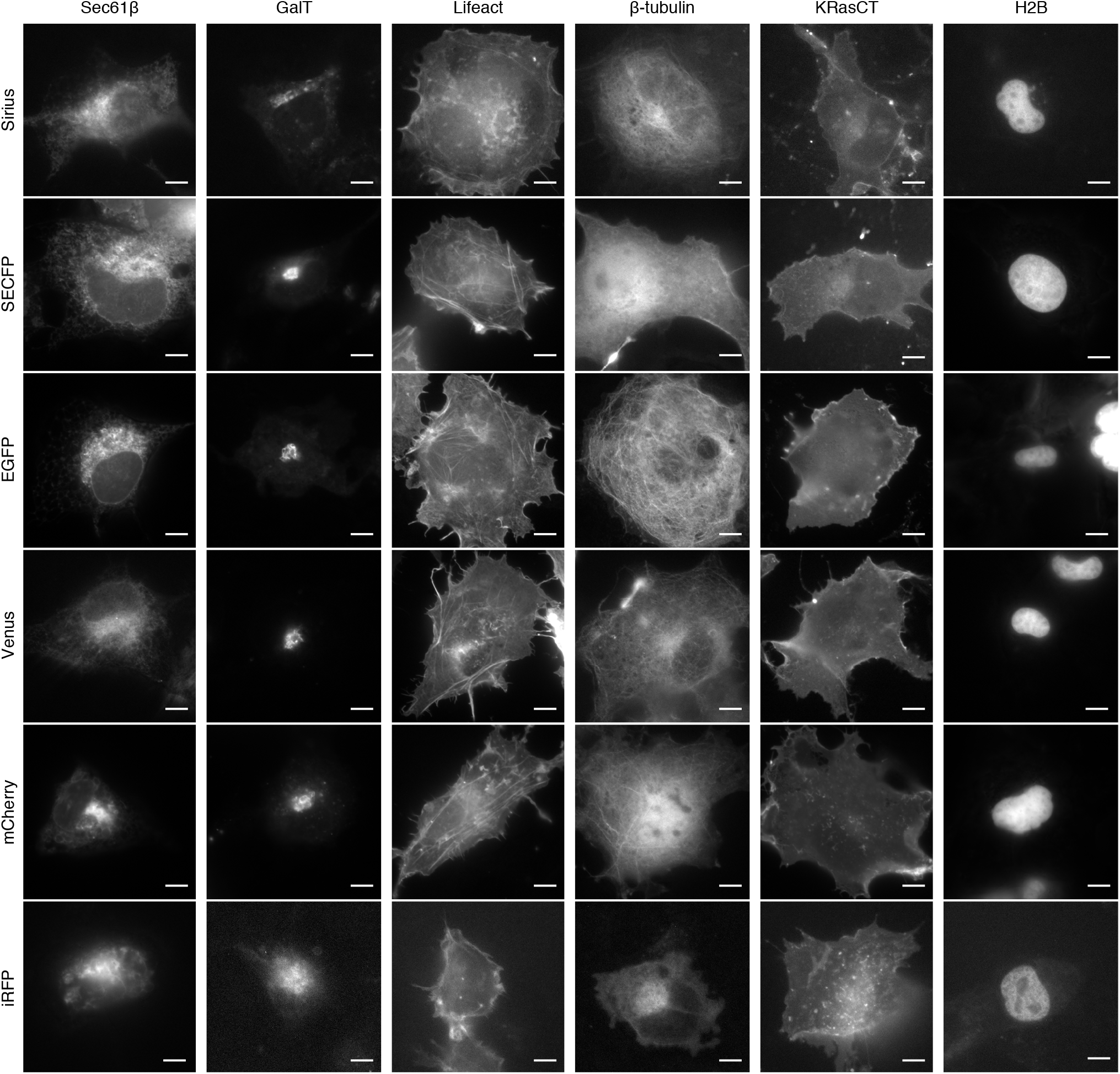
A color pallet of organelle-targeted FFPs (related to Figure 1). Higher magnification images of the panels in Figure 1 are shown. Bar, 10 μm.

**Figure S2:**
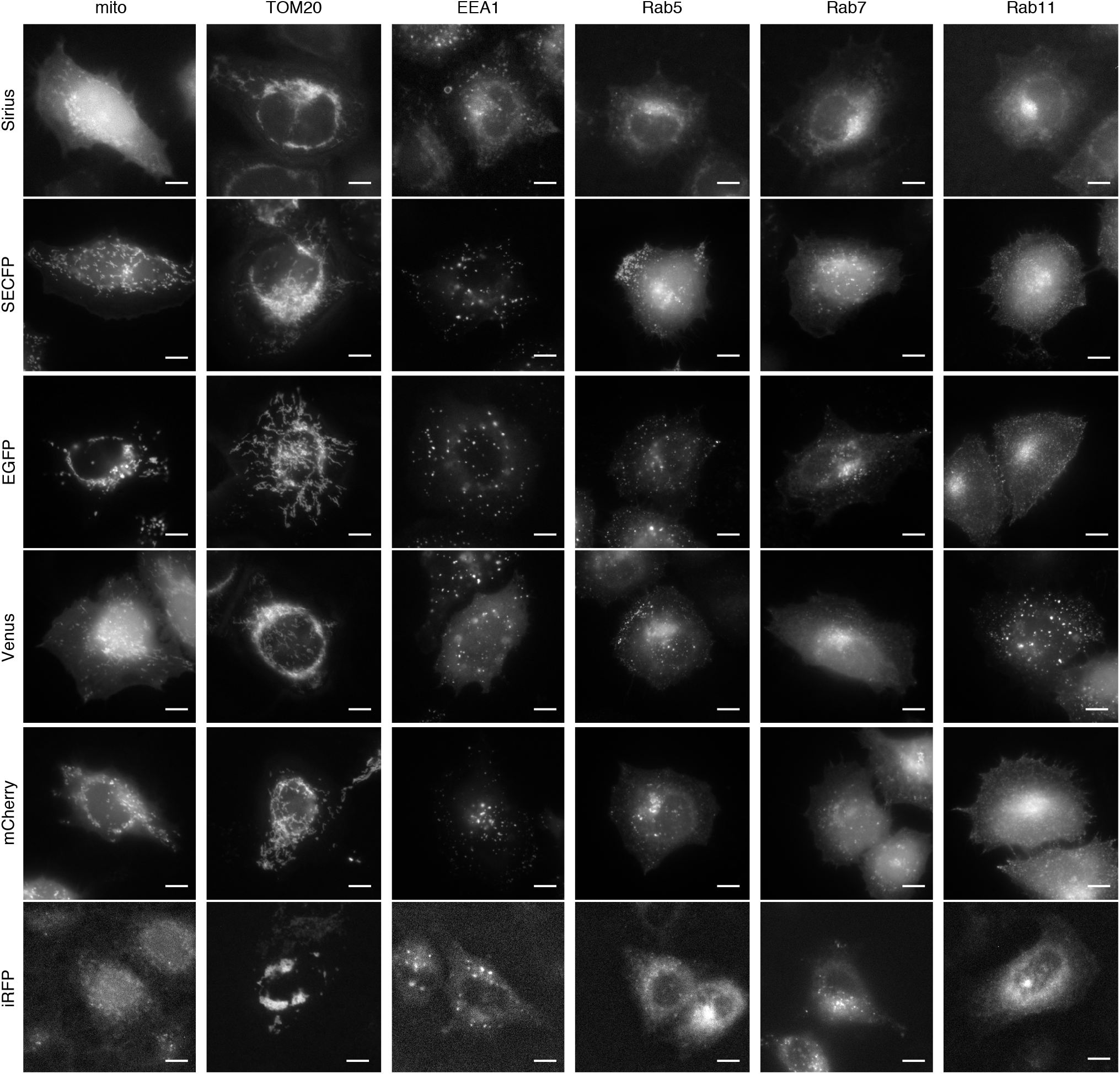
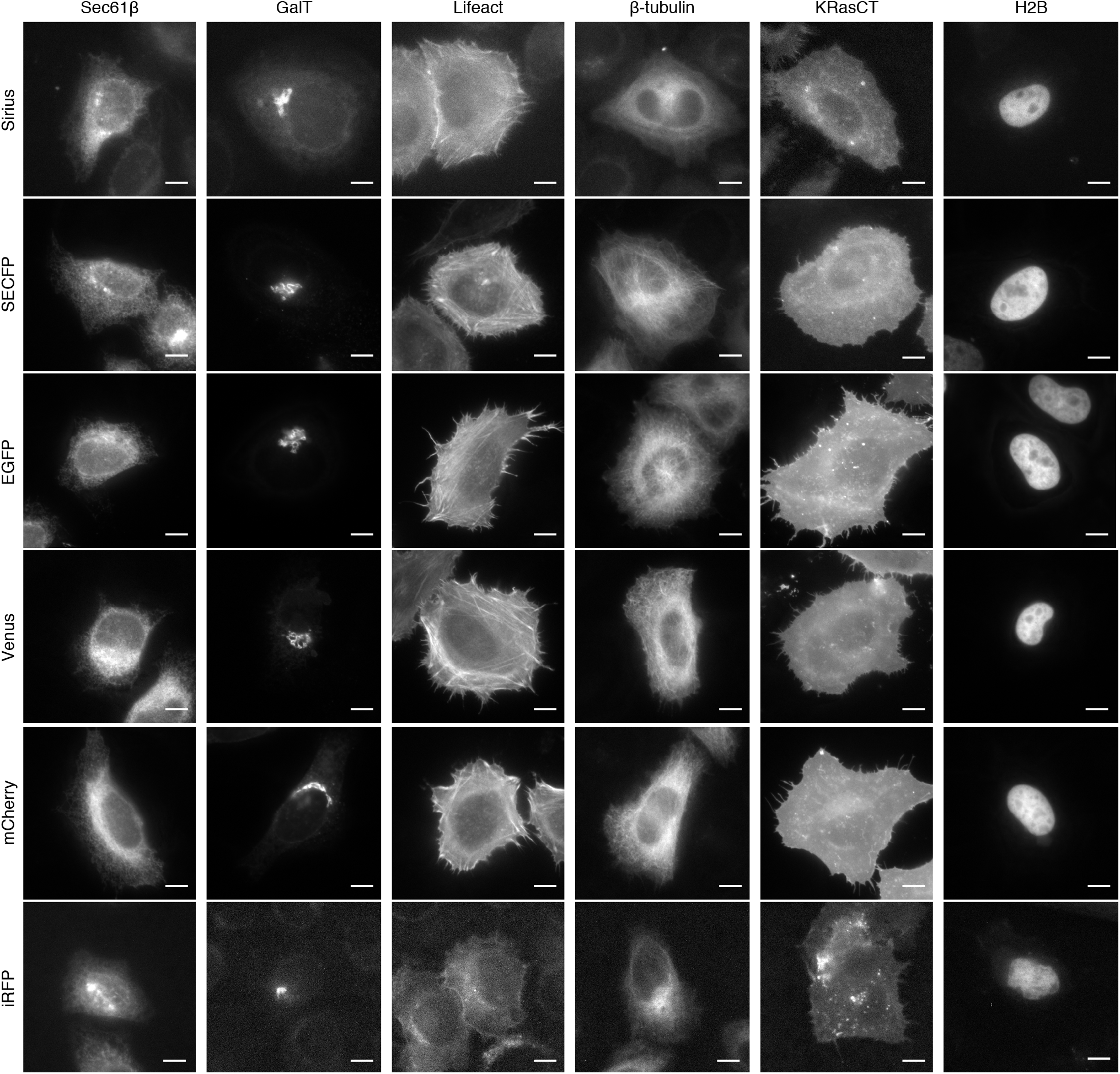
A color pallet of organelle-targeted FFPs (related to Figure 1). HeLa cells were transfected with expression vectors for FFPs (organelle targeted and FP used are indicated at the top and left, respectively) for 24 h and observed with confocal microscopy. Representative images are shown. Bar, 10 μm.

**Figure S3:**
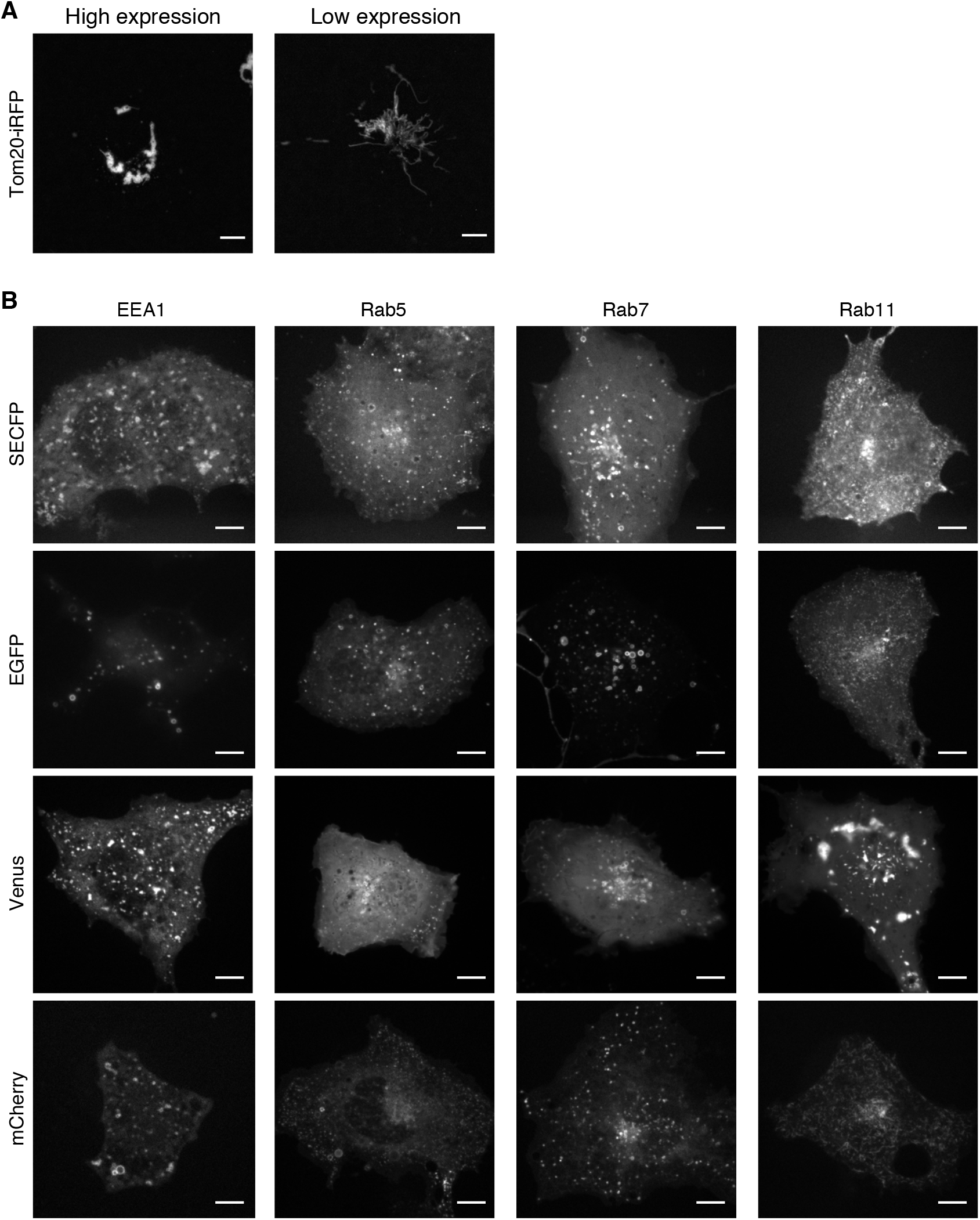
Localization of mitochondria-and endosome-targeted FFPs (related to Figure 1). (**A**) Representative images of Cos-1 cells expressing Tom20-iRFP at higher and lower levels are shown. Bar, 50 µm. (**B**) Cos-1 cells expressing FFPs with organelle-targeting sequences and FPs that are indicated at the top and left, respectively, were observed with confocal microscopy. Representative images are shown. Bar, 10 µm.

**Figure S4:**
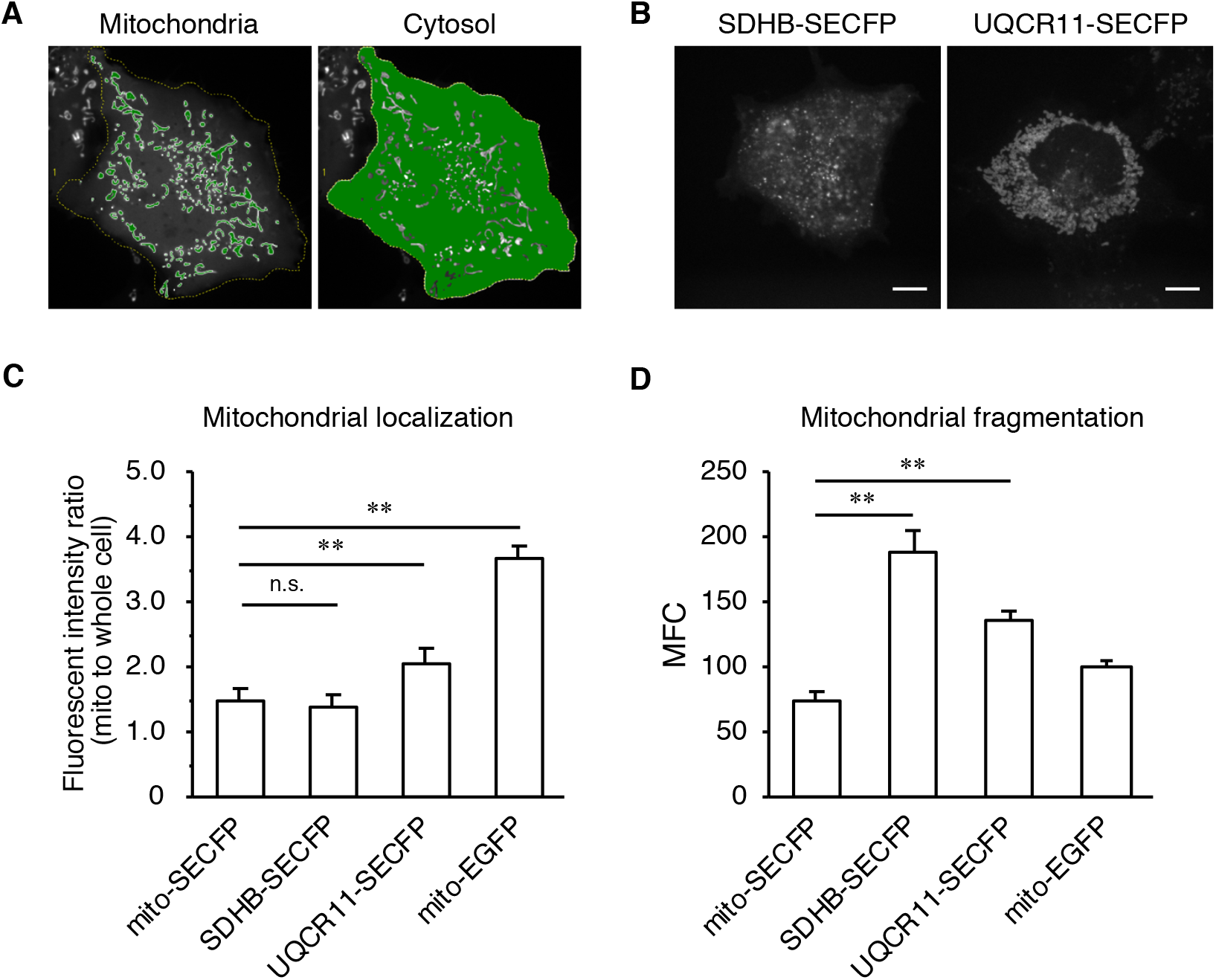
Localization of mitochondria-targeted FFPs with different mitochondrial targeting sequences (related to Figure 1). (**A**) Workflow used for the quantification of the mitochondrial localization of FFPs. (**B**, **C**, **D**) Cos-1 cells transfected with the expression vectors for FFPs indicated at the top were observed with confocal microscopy. Representative images are shown (**B**). The extent of mitochondrial localization was quantitated and plotted (**C**). Data shown represent the mean ± s.e.m. from three independent experiments (*n* > 10) and were analyzed by one-way ANOVA with a post hoc Tukey HSD test. **, *p* < 0.0001, n.s., not significant. (**D**) The extent of mitochondrial fragmentation was quantitated and plotted. Data shown are the mean ± s.e.m. from three independent experiments (*n* > 10). **, *p* < 0.0001, as calculated by one-way ANOVA with a post hoc Tukey HSD test. n.s., not significant.

**Figure S5:**
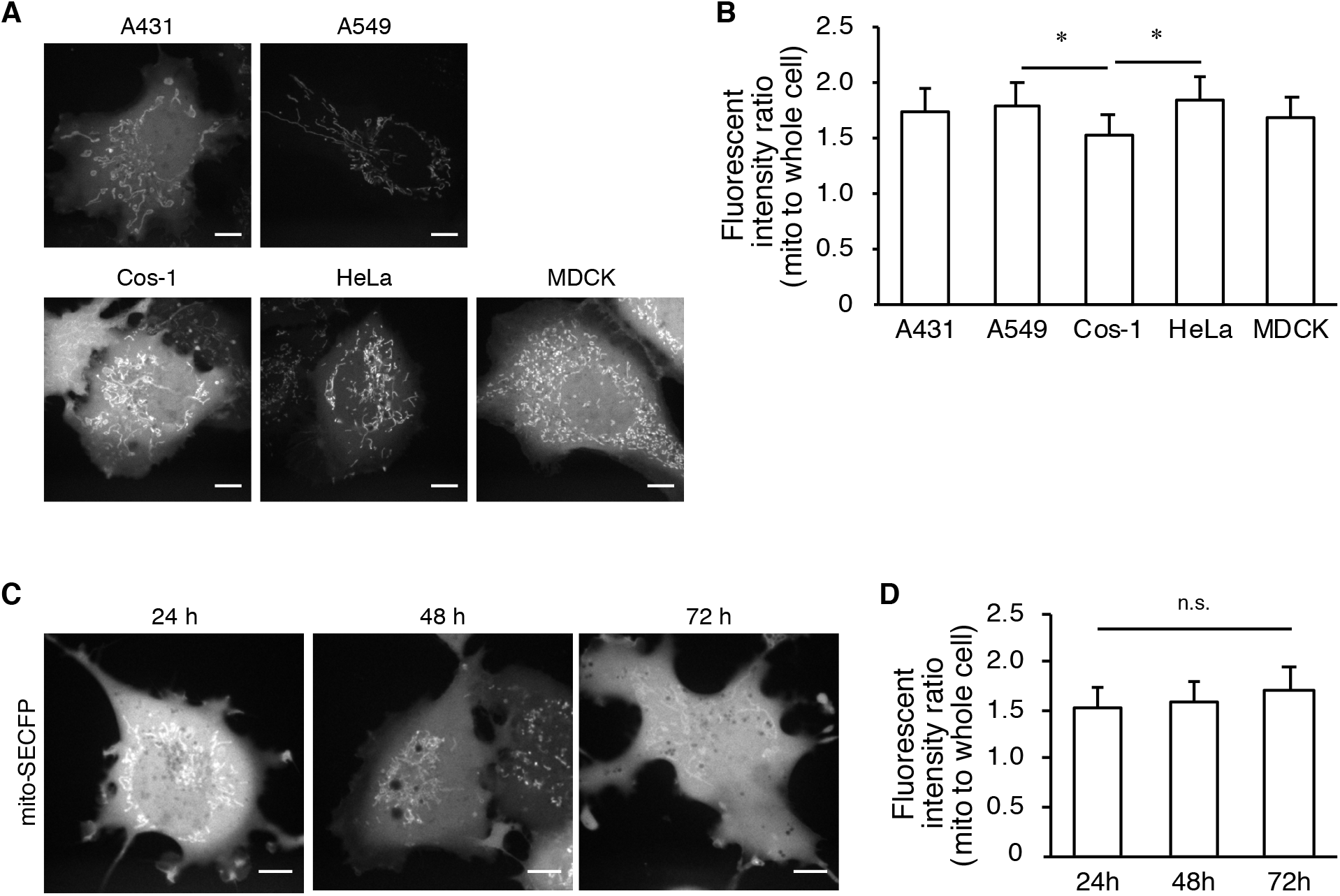
Localization of mitochondria-targeted FFPs in different cell strains or with incubation periods (related to Figure 1). (**A**) Expression vectors for mito-SECFP were transfected into the cell lines indicated at the top. The cells were observed with confocal microscopy. Representative images are shown. Bar, 10 μm. (**B**) The extent of the mitochondrial localization of FFPs was calculated and plotted. Data shown are the mean ± s.e.m. from three independent experiments (*n* > 10). *, *p* < 0.01, as calculated by one-way ANOVA with a post hoc Tukey HSD test. (**C**, **D**) Cos-1 cells expressing mito-SECFP were observed with confocal microscopy at 24 h, 48, and 72 h after transfection. Representative images are shown (**C**). The extent of mitochondrial localization was quantitated and plotted (**D**). Data shown represent the mean ± s.e.m. from three independent experiments (*n* > 10) and were analyzed by one-way ANOVA. n.s., not significant.

**Figure S6:**
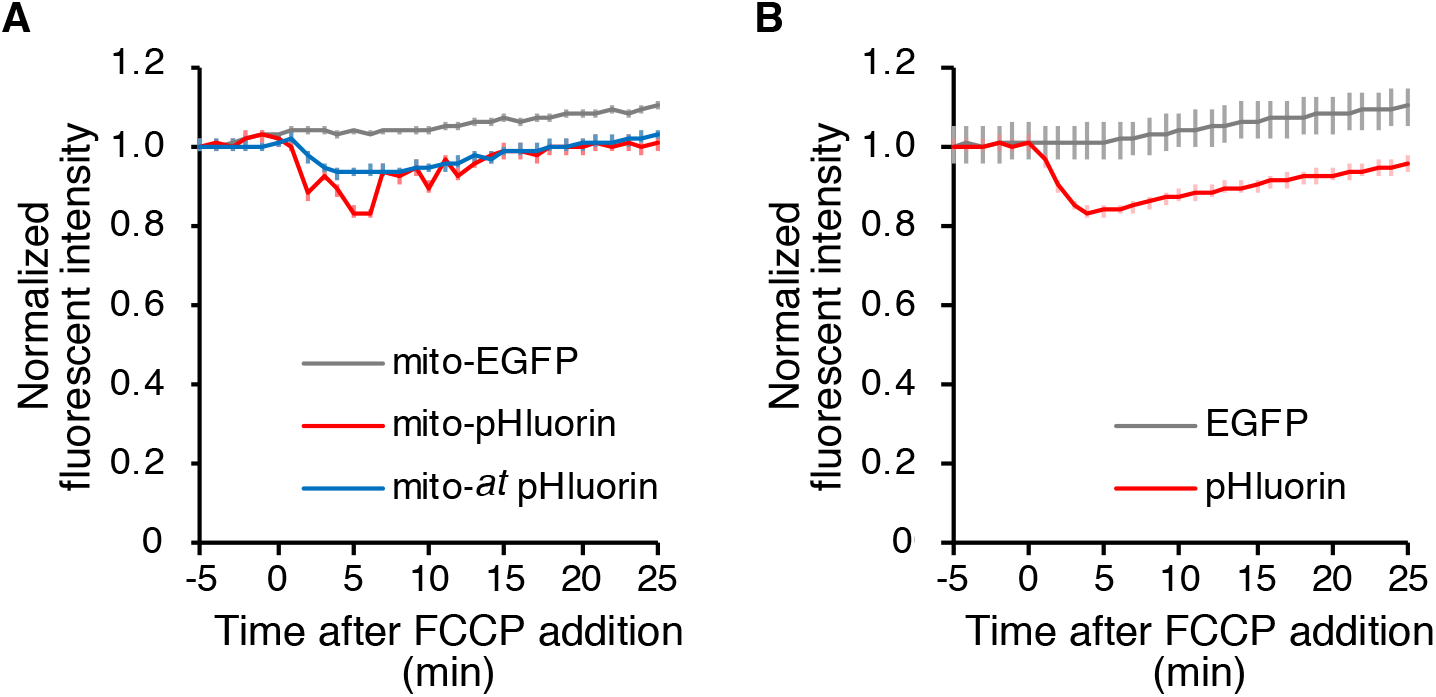
Measurement of mitochondrial pH by FFPs (related to Figure 6). (**A**) Fluorescence intensity in the mitochondrial area was separately quantified using the mitochondria-masked images from Figure 6. (**B**) Cos-1 cells expressing the proteins indicated at the bottom were subjected to time-lapse fluorescence microscopy. At time 0, the cells were exposed to FCCP. The normalized fluorescence intensities in the entire cell area were plotted over time. The data shown are the mean ± s.e.m. from three independent experiments (*n* > 60). *p*-values were calculated by MANOVA. **, *p* < 0.0001, n.s., not significant.

**Supplementary Table 1.**
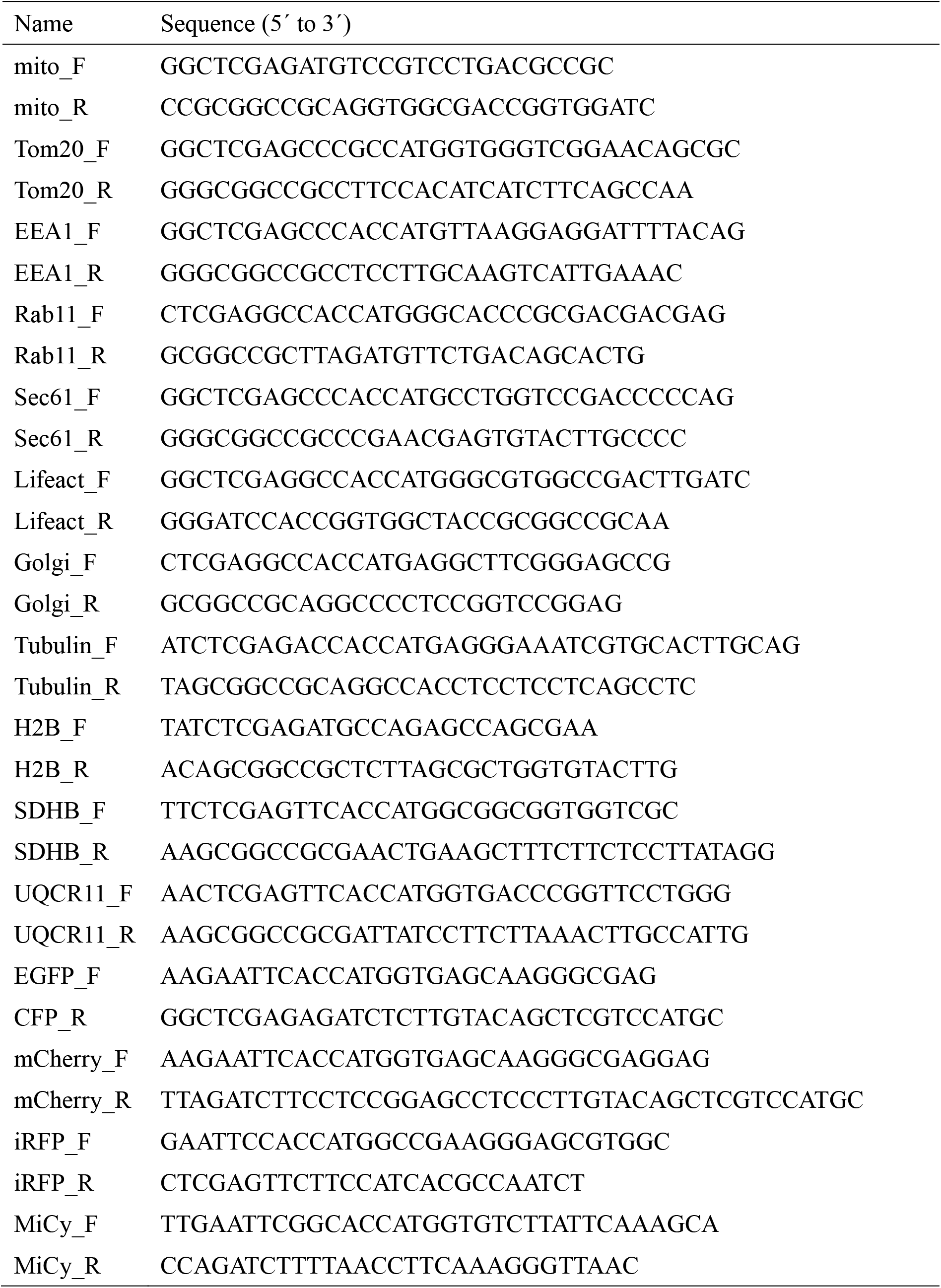
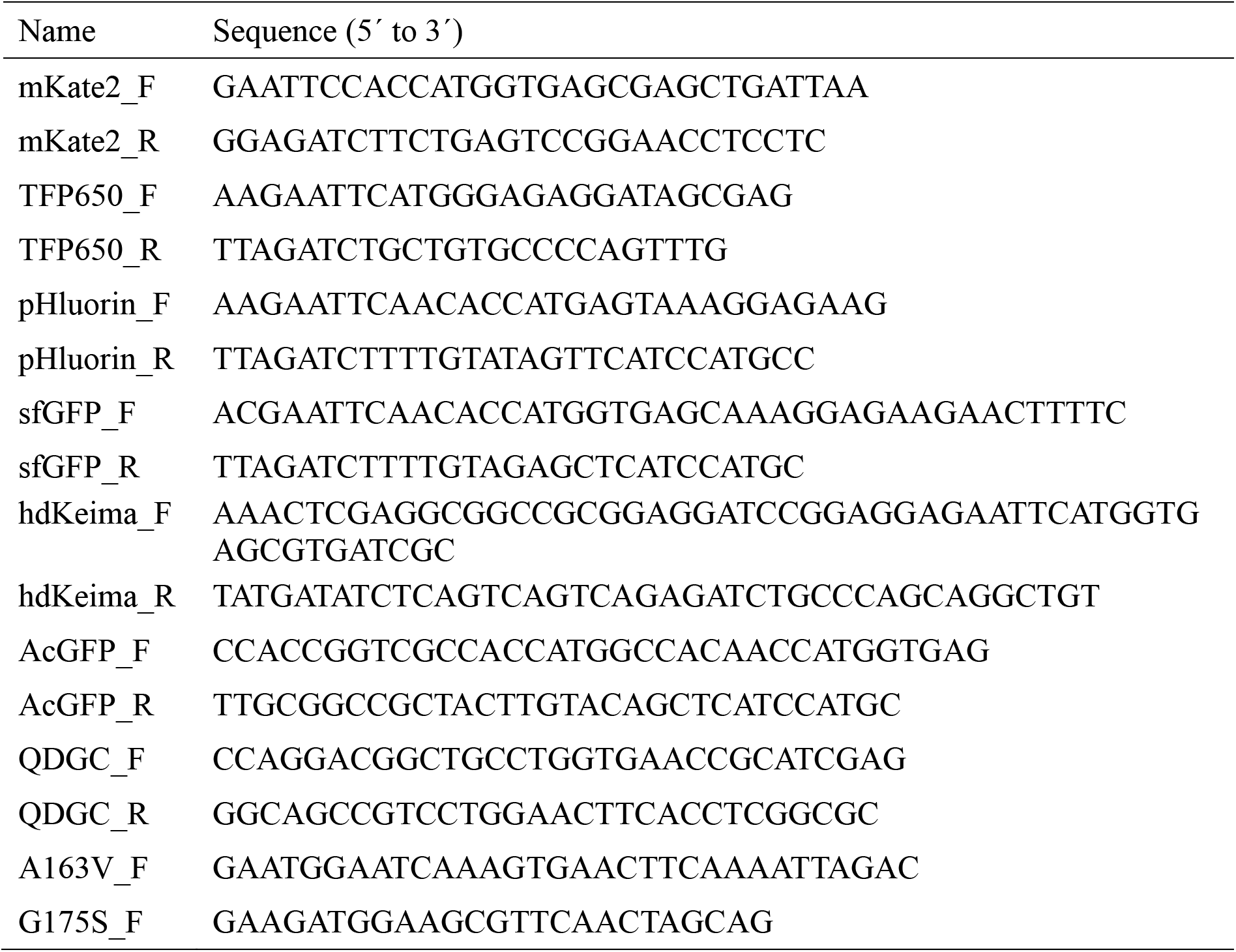
Primers used in this study. Name Sequence (5’ to 3’)

